# The intensity of supplementary feeding in an urban environment impacts overwintering Mallards *Anas platyrchynhos* as wintering conditions get harsher

**DOI:** 10.1101/2024.02.07.579312

**Authors:** Marta Witkowska, Wojciech Wesołowski, Martyna Markiewicz, Jonasz Pakizer, Julia Neumann, Agnieszka Ożarowska, Włodzimierz Meissner

## Abstract

Although urbanization poses various threats to avifauna, some bird species, including Mallards, are attracted to towns and cities as their winter habitats due to favourable temperatures and abundant anthropogenic food. In this study, we investigated how population dynamics changed in relation to winter harshness and intensity of supplementary feeding. The results indicated that the number of Mallards increased with the feeding intensity but the pattern differed in response to worsening of wintering conditions on water bodies with distinct supplementary feeding intensity. The ponds with low feeding intensity were occupied by fewer but more stable populations, suggesting the presence of natural food resources there, available for small number of individuals, regardless of winter conditions. On the ponds with medium feeding intensity, we observed a bell-shaped relationship between the number of ducks and ice cover. Unpredictable anthropogenic food on those ponds might not provide sufficient food resources, forcing birds to relocate when wintering conditions worsen. In such conditions, the ponds with a high feeding intensity attracted more ducks. We demonstrated that the wintering population of Mallards in urban areas is not solely influenced by ambient temperature but is also significantly affected by the intensity of supplementary feeding.

## 1. INTRODUCTION

The ongoing urbanization of the world, with the growing number and areas of cities, as well as the size of human population living in them, are factors rapidly influencing the natural environment and populations of animals. Usually, this impact is reported as negative, since urbanization results in the degradation or loss of habitats and extinction of species, leading to loss of biodiversity (Aronson et al., 2014; Clergeau et al., 2006; Wang et al., 2020; Zhu et al., 2020). Birds within urbanized areas are at risk of mortality due to collisions with built structures (Doren et al., 2021), the presence of domesticated predators (Loss and Marra, 2017; Pavisse et al., 2019), and increased transmission of potentially pathogenic microorganisms (Meissner et al., 2015a). Moreover, cities are noisier than rural areas and that limits birds’ vocal communication (Brumm, 2004; Slabbekoorn and Ripmeester, 2008), while elevated artificial light pollution causes shifts in their daily and annual cycle events (Dominoni, 2015). Yet, at the same time, cities provide a new type of environment, that may be utilized by some species as a living habitat, either by colonizing the urban spaces and constructions, or using green spaces inside cities, that are intended to welcome wildlife and at the same time allow citizens to interact with it (Chace and Walsh, 2006; González-Lagos et al., 2021). Such interaction was reported to be beneficial for people living in the cities, with supplementary feeding of animals being one of the most commonly practiced activities of this kind, which brings citizens into closer contact with wildlife (Rowan and Beck 1994; Cox and Gaston, 2016). Moreover, higher winter temperatures in cities than surrounding rural areas (Arnfield, 2003; Collier, 2006) diminish thermoregulatory costs and result in decreased migratory tendencies (Bonnet-Lebrun et al., 2020). As food resources availability is a key factor determining animal population structure and distribution, the provision of anthropogenic food both intentional (e.g., feeders in private gardens, feeding wildlife in parks and public gardens) and unintentional (e.g., trashes with leftover food, remains from feeding domesticated animals) may locally improve feeding conditions and therefore increase the population size and density (Cox and Gaston, 2018; Fuller et al., 2008; Plummer et al., 2019).

The Mallard *Anas platyrhynchos* is a dabbling duck species, that has adapted to the urban environment, and extensively uses the waterbodies located there during different stages of its annual cycle (Jarman et al., 2020; Kopij, 2020; Meissner et al., 2015b). For overwintering individuals, cities provide higher ambient winter temperatures (Kalnay and Cai, 2003), keeping the surfaces of the waterbodies from freezing, and therefore allowing Mallards to obtain their natural food. This, together with the supplementary feeding provided by citizens may add up to abundant food resources offered by the urban environment. Additionally, higher ambient temperatures within cities may create fewer constraints on keeping the metabolic rate essential for maintaining the stable body temperature of endothermic birds, that finally lowers the costs of maintenance of individuals overwintering in the cities, compared to their conspecifics wintering in more natural habitats (Tryjanowski et al., 2015). The importance of ambient temperature changes and winter harshness were pointed out as main factors influencing the demographic parameters of the Mallard population either in the urban or rural environment, such as its size (Meissner et al., 2015b; Schummer et al., 2010), sex-ratio (Meissner and Witkowska, 2023; Pattenden and Boag, 1989), and survival (Bergan and Smith, 1993; Manikowska-Slepowrońska and Meissner, 2022).

In urbanized areas, Mallards gather in large numbers in sites where people feed them (Avilova and Eremkin, 2001; Engler et al., 1988). Yet, the studies of urban overwintering waterbirds usually only speculate about the importance of supplementary feeding and its consequences for the population, without including it as a factor in the research design (Avilova and Eremkin, 2001; Meissner and Witkowska, 2023) but see: Polakowski et al., 2010). A number of conducted research support the existence of supplementary feeding consequences during winter, but mostly for land birds using garden feeders, particularly passerines, and the increase in population density is one of the effects reported in those studies (e.g., Fuller et al., 2008). Moreover, supplementary feeding may contribute to species range expansion (Greig et al., 2017) and diversity of bird communities (Plummer et al., 2019). In case of Mallards wintering in urban area, supplementary feeding contributes to higher survival of females during the periods of low temperatures despite increased energy expenditure (Meissner and Witkowska, 2023). Thus, we suspect that the changes in the abundance of the Mallard populations wintering in urban areas might be a response to the worsening of wintering conditions, which may have a different outcome depending on the intensity of anthropogenic food provision, pronounced even at the local scale. Therefore, this study aimed to describe the changes in the abundance of Mallards wintering in urban area in relation to the decrease in the ambient temperature and freezing of the ponds occupied by this species during winter, taking into account the differences in the intensity of feeding conducted by the citizens on a given pond.

## 2. METHODS

### 2.1 Fieldwork

The survey has been conducted on fifteen different ponds located on the Oliwski Stream in Gdańsk (Poland 54°21′N, 18°40′E; Fig. 1) for five consecutive winter seasons since 2017. We counted female and male Mallards separately, every two weekends, starting from the third weekend of December, until the second weekend of March, which resulted in seven counts each season. This protocol was not followed during the first, pilot season of winter 2017/2018, where we counted birds on five occasions only, from the last weekend of January to the second weekend of March (Table 1S). Nonetheless, we included the data collected during that season in our analysis. Each counting event started at 8:00 AM in the same place, being the central pond in the middle of Oliwski Stream, followed by one person conducting the counting on seven ponds located upstream and the other person surveying eight ponds located downstream from the starting point. The total time of the census did not exceed three hours.

**Table 1.**
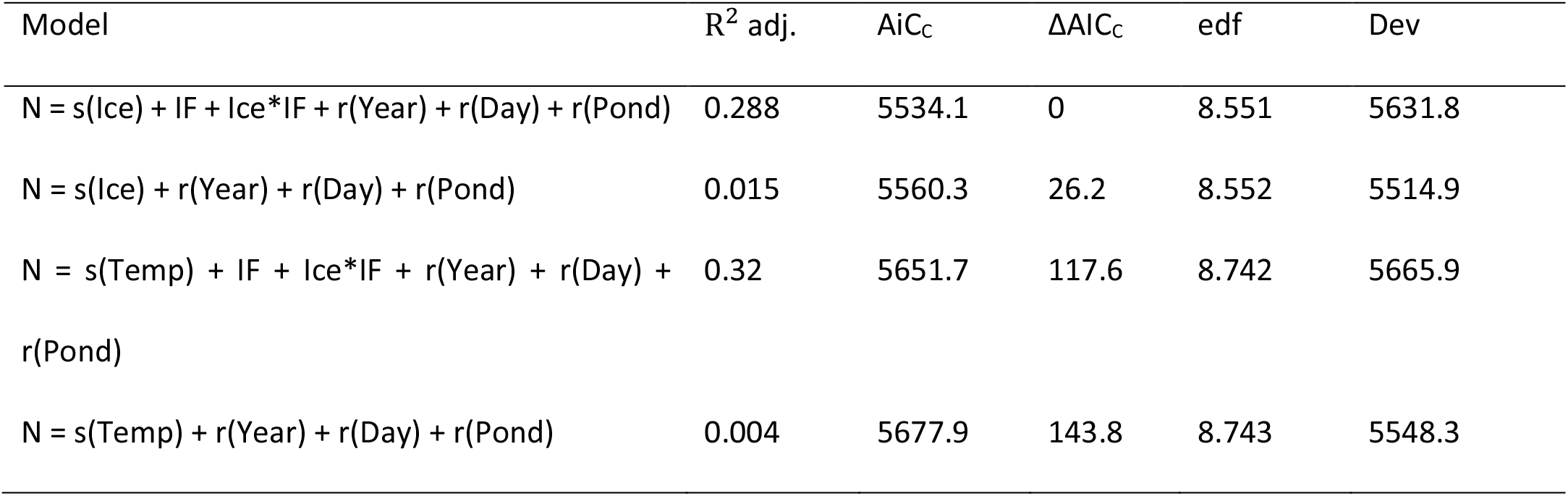
Ranking of generalized additive mixed models and generalized linear mixed models used to estimate the number of wintering Mallards (N). Model factors included: Ice – ice cover on a given pond [%]; Temp – ambient temperature [°C]; IF – category of the intensity of anthropogenic feeding on a given pond; Year - season, Day – consecutive counting corresponding to the same weekend of a given month in different seasons; Pond – given pond; s() – the smooth term; r() – random factor; * - interaction between independent variables; R^2^adj. – adjusted R^2^, AIC_C_ - Akaike’s Information Criterion; ΔAIC_C_ – the difference between AIC_C_ of a given model and the model with the lowest AIC_C_; edf – effective degrees of freedom; Dev – deviance.

**Fig. 1.**
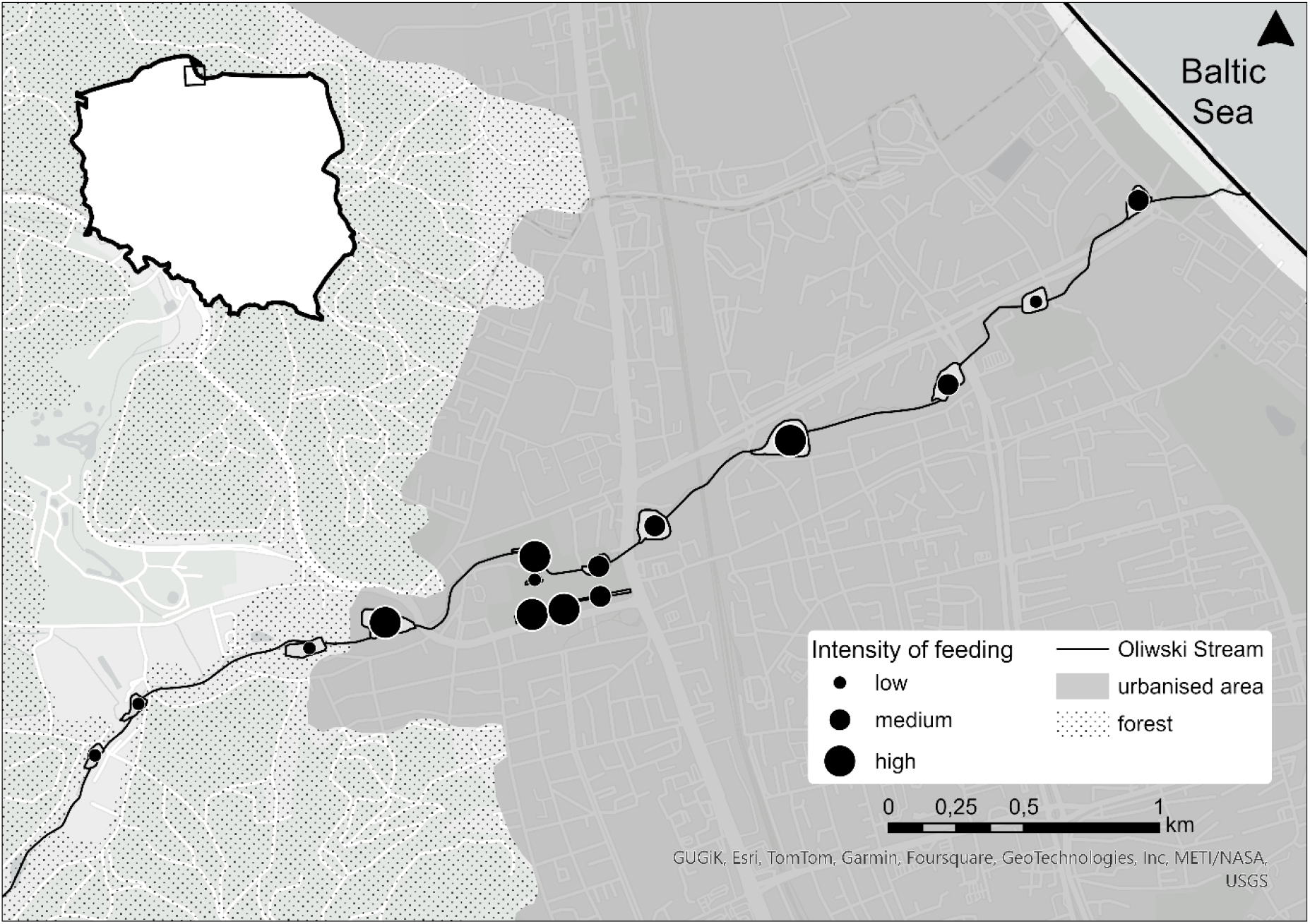
Location of 15 ponds on the Oliwski Stream in Gdańsk.

To establish the intensity of anthropogenic feeding conducted on each pond (later referred to as ‘IF’) we conducted the observations of citizens feeding Mallards and other waterbirds (i.e., gulls, mute swans *Cygnus olor*, and other species of ducks) occupying a given pond during the weekend, and we counted the incidents of feeding birds. We considered 9:00 AM till 3:00 PM to be rush hours for citizens’ outdoor activities, and within that time frame we established eight, 45-minute-long intervals, in which we conducted our observations. In each interval, a given pond was observed for 30 minutes and the rest of the remaining time was used by the observer to commute from one pond to another. To create a sequence of ponds visited by an observer in consecutive time intervals, in a way that ensured collecting representative data and minimizing any time-related bias, we picked the ponds to observe at random.

### 2.2 Environmental conditions

Along with the census of Mallards, we estimated the percentage of ice cover on the surface of the pond. Ambient temperature was calculated as a mean temperature from 5 days preceding the date of the survey, as the response of Mallards to a drop in temperature does not have to be immediate and short-term chilling might not promote their movements (Sauter et al., 2010). The data on ambient temperature was taken from the reports of the meteorological station located at Gdańsk Rębiechowo Airport, i.e., approximately 7 km away from the studied site. The winter season in the studied area is rather mild, as the mean winter temperature from five years of the study was 1.5 °C, and in the case of only five counting events the mean ambient temperature during the day was below 0 °C (Fig. 2, Table 1S).

**Fig. 2.**
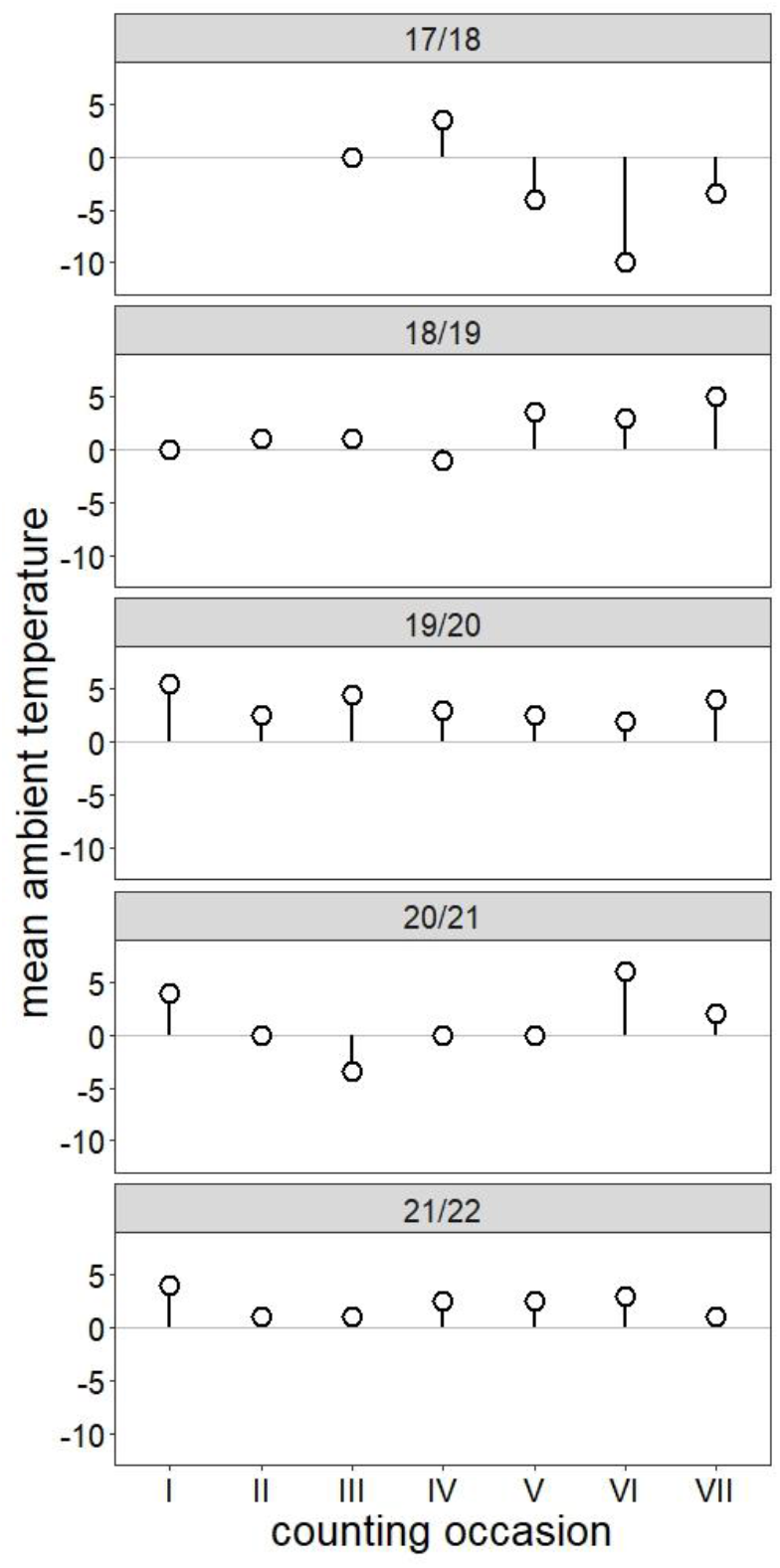
Mean ambient temperature established for a given counting occasion in 5 consecutive seasons.

### 2.3 Statistical analysis

We established three different categories of IF, by taking the distribution of the total number of feeding incidents conducted on the particular pond and dividing them using the breakpoints at 33% and 67% of the distribution range. With this approach, five ponds were categorized with low IF, five ponds with medium IF, and five ponds with high IF (Fig. 1). To check how the number of Mallards was influenced by temperature and ice cover on ponds with different categories of IF, we used the Generalized Additive Mixed Model (GAMM) with Poisson distribution and log-link function (Hastie and Tibshirani, 1986), computed with *gmm4* package in R 4.2.2 (R Core Team, 2021; Wood et al., 2020), where IF categories were set as a fixed parameter, and either temperature or ice coverage (due to strong collinearity between those two parameters: Pearson correlation r = 0.85, p < 0.001) as smoothing term. We considered both additive and interactive relationships between those two covariates. Year, a sequence number of consecutive counting corresponding to the same weekend of a given month in different seasons, and the pond were established as random parameters to control for their randomized effects. We used the Akaike Information Criterion for small sample size (AIC_c_) and adjusted R^2^ to rank the models, considering the model with the lowest AIC_c_ and the highest adjusted R^2^ to be the most informative (Burnham and Anderson, 2004)

## 3. RESULTS

According to the best-fitting model the number of wintering Mallards was influenced by the extent of ice cover along with intensity of anthropogenic feeding (IF) and an interaction between those covariates while controlling for all random factors such as season, date of the census, and pond. This model was also strongly supported by the data compared to the next models (Table 1). All fixed factors significantly affected the number of wintering Mallards. Although the inclusion of random effects increased the fit of models, the percentage of variance explained by them was low (Table 2). The models including the percentage of ice cover instead of ambient temperature had higher support as well as the models that included IF together with an environmental condition (either ambient temperature or ice coverage) as a fixed factor (Table 1).

**Table 2.**
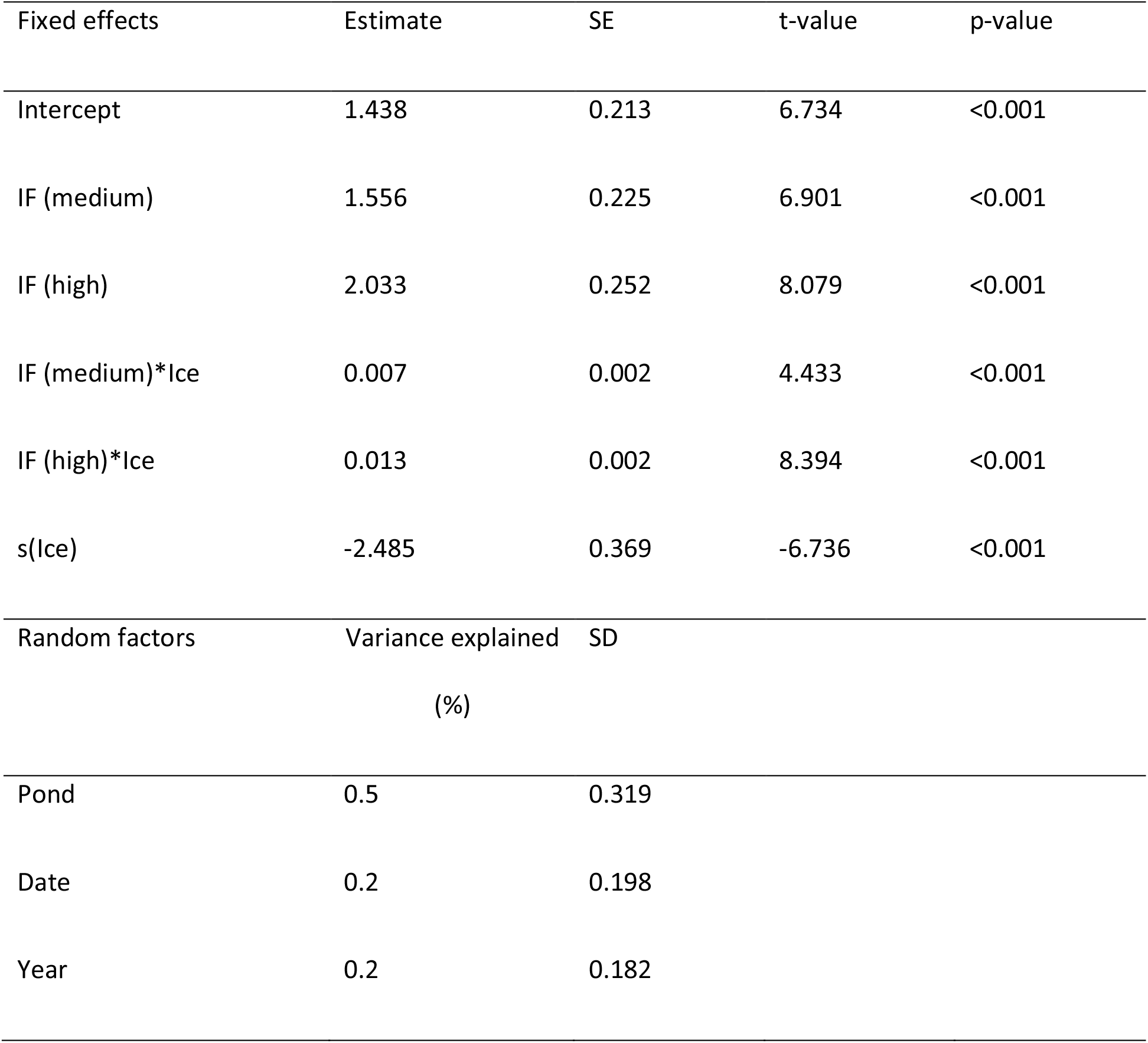
Estimates of the Generalized Additive Mixed Model for the effects of the best-fitted model for the number of wintering Mallards.

The number of wintering Mallards varied between the ponds characterized by different IF categories (Poisson GAMM; χ^2^ = 81.962; p < 0.001). The highest number of wintering Mallards was noted on the ponds with high IF and the smallest on ponds with low IF, with medium numbers of wintering Mallards on ponds with medium IF (Fig. 3, Table 2).

**Fig. 3.**
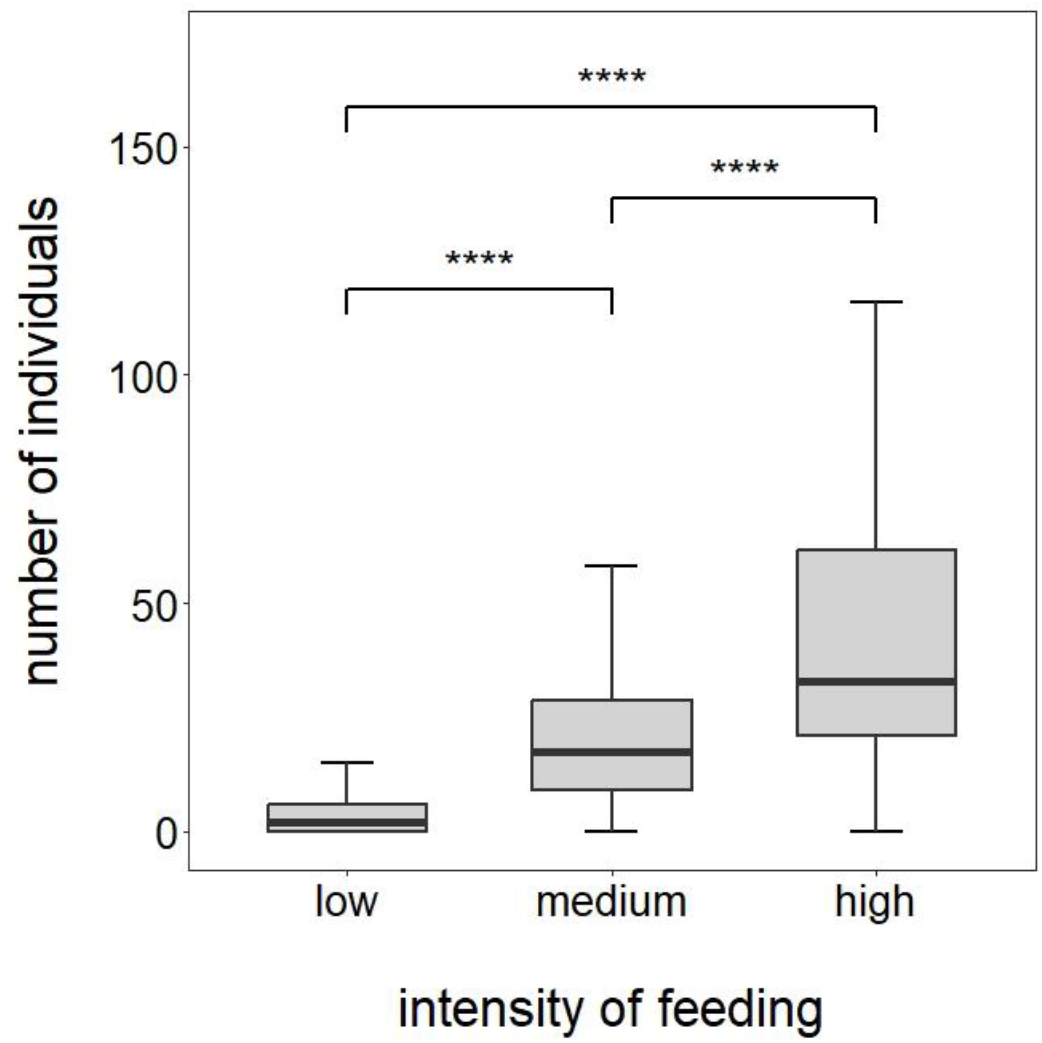
Number of wintering Mallards on the ponds with different IF. Statistically significant differences according to the Generalized Additive Mixed Model are given, where ns – no statistically significant differences; **** - p < 0.0001

On the ponds with low IF the number of wintering Mallards decreased with the increase in ice cover. This relationship was a nonlinear on medium IF ponds, i.e., initially the number of Mallards occupying such sites had increased, but when the ice cover reached approximately 50% of the pond’s surface, the number of ducks started to decrease. The ponds with high IF were characterized by an increase in the number of wintering Mallards despite growing ice cover on the surface of these water bodies (Fig. 4).

**Fig 4.**
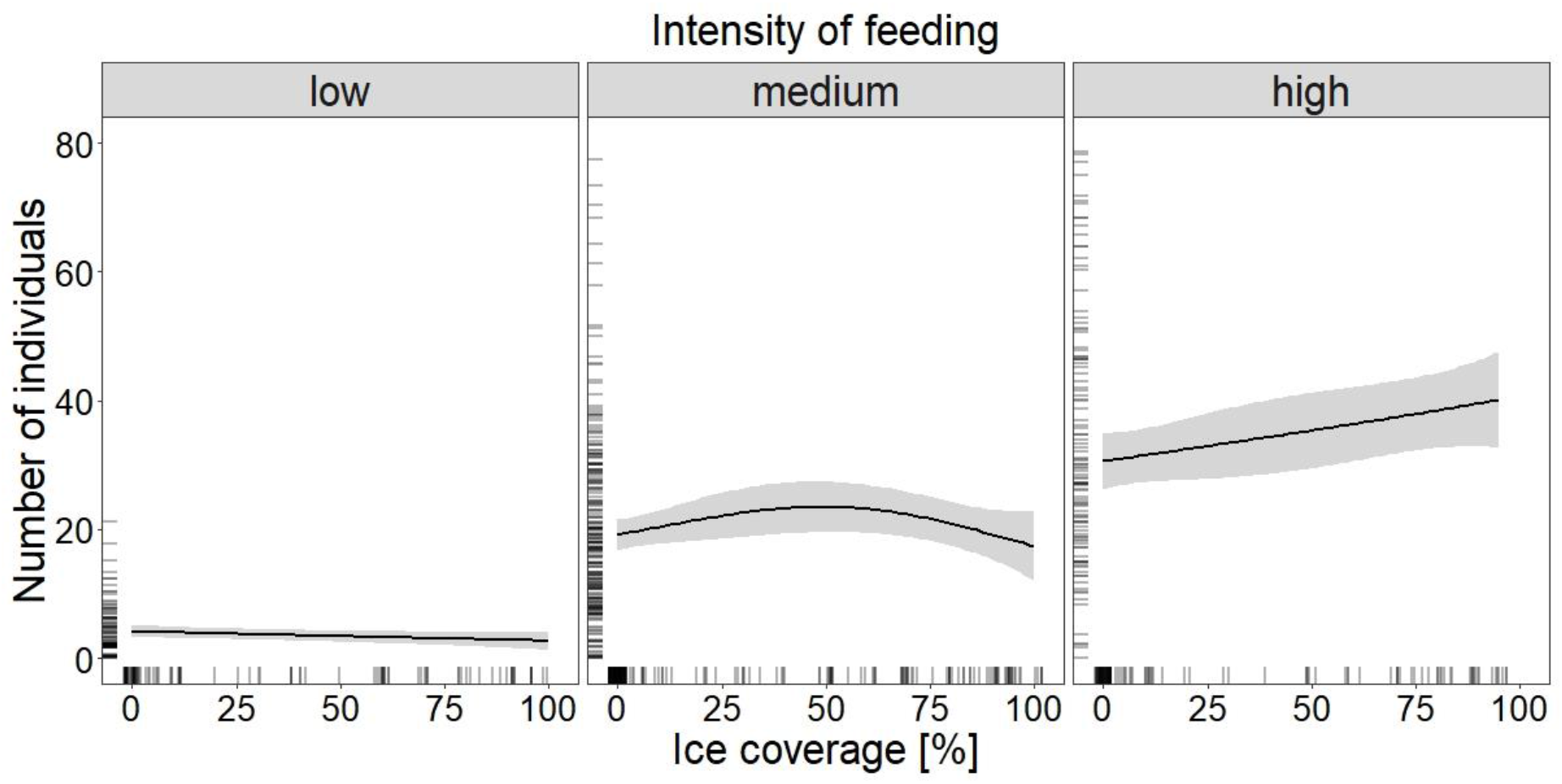
Relationship between ice cover of the surface of a given pond and the number of wintering Mallards on the ponds with three different IF. The black line – significant relationship estimated with the Generalized Additive Mixed Model; grey shaded area – 95% confidence interval. Bars at the bottom and left-hand side of the graph indicate a presence of a sample.

## 4. DISCUSSION

Despite several factors increasing bird mortality in towns and cities (Doren et al., 2021; Meissner et al., 2015a; Pavisse et al., 2019), some species tend to select urban areas as their wintering habitats, due to higher and more stable ambient temperature and abundant anthropogenic food resources (Luniak 2004; Tryjanowski et al., 2015). Both these factors are important in maintaining a suitable metabolic rate for the survival of an individual, which may be compromised, especially in harsh wintering conditions, when natural food becomes scarce and cold weather increases the energetic demand of endothermic animals. Supplementary feeding in urban areas was previously accounted for causing flocking of different bird species, including wintering Mallards (Fuller et al., 2008; Plummer et al., 2019; Polakowski et al., 2010). In this study, we provide more detailed information on how the intensity of this activity affects wintering Mallard community in relation to winter harshness. In the proposed set of models, it was winter harshness defined as ice coverage of a given pond that was more important factor than mean daily ambient temperatures. Still, this might result from the fact that we analyzed the temperatures measured only in one meteorological station, fairly distant from the studied ponds, so this approach failed to give detailed information on the mean daily temperature characteristic for each of the ponds. On the other hand, the ice cover of a given pond is a local indicator of the severity of overwintering conditions as it limits the pond’s area available for ducks and therefore is a better predictor for the abundance of wintering Mallards based on the proposed models. The number of individuals generally increased with the growing intensity of feeding reported for a given pond, which shows that an additional source of food allows for larger communities of overwintering Mallards to gather in a given area. Such a result has been previously reported from another Polish city (Polakowski et al., 2010). In this study, in addition to previous findings, we documented different patterns of population number dynamics in relation to both intensity of supplementary feeding and worsening of wintering conditions, i.e., freezing of waterbodies that limits open water surface area. On the ponds with low intensity of feeding, the number of overwintering ducks was the lowest and most stable in regard to lowering ambient temperatures and growing ice cover of the ponds’ surface. Such ponds and their surroundings may probably offer some natural food resources, suitable for a low number of individuals, that surprisingly stayed there even when the unfrozen water surface was strongly limited. Wintering Mallards to some extent can endure harsh winter conditions by lowering their activity and consequently limiting their energy expenditure and food intake, by waiting out the complete freeze over of waterbodies (Meissner and Markowska, 2009). In continental scale, wintering Mallards more than other duck species resist freezing temperatures if the waterbody is not completely covered with ice (Sauter et al., 2010; Schummer et al., 2010). Perhaps such strategy is more profitable compared to moving to different wintering sites, but in the ponds with low feeding intensity it may be utilized only by a small number of individuals. Medium intensity of feeding enhances the population abundance, but with the deterioration of wintering conditions, the carrying capacity of the environment with such supplementation of food is too low to maintain favorable conditions for a large number of individuals. Moreover, lowering temperatures combined with freezing of waterbodies, increase the energetic requirements to maintain stable body temperatures of endothermic birds (Swanson and Olmstead, 1999). In such cases natural food becomes scarce and anthropogenic food turns into an insufficient and unpredictable resource causing birds occupying such ponds to abandon these suboptimal habitats. It is not known whether those individuals moved to the ponds with high intensity of feeding, still we observed constant increase in the number of Mallards on those ponds with worsening of wintering conditions. This may be supported by previous studies based on the ring recovery analyses indicating that some individuals can move within an urban area towards the sites with more intensive supplementary feeding (Polakowski et al., 2010). It is possible that such short-distance movements occur prior to longer migratory movements that were reported in Mallards as a response to significant increase in severity of winter (Sauter et al., 2010). However, it is worth mentioning, that in five years of conducting this project, only five counting events were carried out in the conditions when mean daily ambient temperature was below 0°C. Therefore, we assume that based on the collected data we were not able to demonstrate a situation of truly harsh winter in which further deterioration of wintering conditions would probably force most birds to leave the area, even of the ponds with the high intensity of feeding.

Mallards wintering in urban areas react to local temperature changes by altering their abundance (Meissner et al., 2015b) and sex ratio (Meissner and Witkowska, 2023). In this study, we also showed that ambient temperature and consequently ice cover had a significant influence on the number of Mallards wintering in the urban area. Yet the response of wintering ducks to these factors reflected by the changes in their numbers differed depending on the intensity of anthropogenic feeding at a particular site. Hence, all these factors should be taken into account in further studies on the abundance of Mallards and other waterbirds wintering in towns and cities.

## ACKNOWLEDGMENTS

We are grateful to all members and sympathizers of the Student’s Scientific Club KOS that took part in the fieldwork.

## DATA AVAILABILITY

Data will be made available on request.

## FUNDGIN

This research did not receive any specific grant from funding agencies in the public, commercial, or not-for-profit sectors

## AUTHORS CONTRIBUTIONS

MW: Conceptualization, Data curation, Formal Analysis, Methodology, Visualization, Writing – original draft

WW: Conceptualization, Data curation, Methodology, Writing – review and editing

MM: Data curation, Methodology, Visualization, Writing – review and editing

JP: Data curation, Writing – review and editing

JN: Data curation, Writing – review and editing

WM: Conceptualization, Supervision, Writing – review and editing

AO: Conceptualization, Supervision, Writing – review and editing

## DECLARATION OF COMPETING INTREST

Authors declare no financial and/or personal relationship that could bias this work.

## Notes

### Competing Interest Statement

The authors have declared no competing interest.

## REFERENCES

Arnfield, A.J., 2003. Two decades of urban climate research: A review of turbulence, exchanges of energy and water, and the urban heat island. International Journal of Climatology 23, 1–26. 10.1002/joc.859

Aronson, M.F.J., la Sorte, F.A., Nilon, C.H., Katti, M., Goddard, M.A., Lepczyk, C.A., Warren, P.S., Williams, N.S.G., Cilliers, S., Clarkson, B., Dobbs, C., Dolan, R., Hedblom, M., Klotz, S., Kooijmans, J.L., Kühn, I., Macgregor-Fors, I., Mcdonnell, M., Mörtberg, U., Pyšek, P., Siebert, S., Sushinsky, J., Werner, P., Winter, M., 2014. A global analysis of the impacts of urbanization on bird and plant diversity reveals key anthropogenic drivers. Proceedings of the Royal Society B: Biological Sciences 281. 10.1098/rspb.2013.3330

Avilova, K. V., Eremkin, G.S., 2001. Waterfowl wintering in Moscow (1985-1999): Dependence on air temperatures and the prosperity of the human population. Acta Ornithol 36, 65–71. 10.3161/068.036.0101

Bonnet-Lebrun, A.S., Manica, A., Rodrigues, A.S.L., 2020. Effects of urbanization on bird migration. Biol Conserv 244. 10.1016/j.biocon.2020.108423

Brumm, H., 2004. The impact of environmental noise on song amplitude in a territorial bird. Journal of Animal Ecology 73, 434–440. 10.1111/j.0021-8790.2004.00814.x

Burnham, K.P., Anderson, D.R., 2004. Multimodel Inference: Understanding AIC and BIC in Model Selection. Sociol Methods Res 33, 261–304.

Chace, J.F., Walsh, J.J., 2006. Urban effects on native avifauna: A review. Landsc Urban Plan 74, 46–69. 10.1016/j.landurbplan.2004.08.007

Clergeau, P., Croci, S., Jokimäki, J., Kaisanlahti-Jokimäki, M.L., Dinetti, M., 2006. Avifauna homogenisation by urbanisation: Analysis at different European latitudes. Biol Conserv 127, 336–344. 10.1016/j.biocon.2005.06.035

Collier, C.G., 2006. The impact of urban areas on weather. Quarterly Journal of the Royal Meteorological Society 132, 1–25. 10.1256/qj.05.199

Cox, D.T.C., Gaston, K.J., 2018. Human–nature interactions and the consequences and drivers of provisioning wildlife. Philosophical Transactions of the Royal Society B: Biological Sciences 373. 10.1098/rstb.2017.0092

Cox, D.T.C., Gaston, K.J., 2016. Urban bird feeding: Connecting people with nature. PLoS One 11, 1–13. 10.1371/journal.pone.0158717

Dominoni, D.M., 2015. The effects of light pollution on biological rhythms of birds: an integrated, mechanistic perspective. J Ornithol 156, 409–418. 10.1007/s10336-015-1196-3

Doren, B.M. van, Willard, D.E., Hennen, M., Horton, K.G., Stuber, E.F., Sheldon, D., Sivakumar, A.H., Wang, J., Farnsworth, A., Winger, B.M., 2021. Drivers of fatal bird collisions in an urban center. PNAS 118, e2101666118. 10.1073/pnas.2101666118/-/DCSupplemental.y

Engler, J., Keller, M., Leszkowicz, J., Zawadzki, J., 1988. Synurbization of the mallard Anas platyrhynchos in Warsaw. Acta Ornithol 24.

Fuller, R.A., Warren, P.H., Armsworth, P.R., Barbosa, O., Gaston, K.J., 2008. Garden bird feeding predicts the structure of urban avian assemblages. Divers Distrib 14, 131–137. 10.1111/j.1472-4642.2007.00439.x

González-Lagos, C., Cardador, L., Sol, D., 2021. Invasion success and tolerance to urbanization in birds. Ecography 44, 1642–1652. 10.1111/ecog.05826

Greig, E.I., Wood, E.M., Bonter, D.N., 2017. Winter range expansion of a hummingbird is associated with urbanization and supplementary feeding. Proceedings of the Royal Society B: Biological Sciences 284. 10.1098/rspb.2017.0256

Hastie, T.J., Tibshirani, R.J., 1986. Generalized additive models. Statistical Science 1, 297–318. 10.1201/9780203753781

Jarman, T.E., Gartrell, B.D., Battley, P.F., 2020. Differences in body composition between urban and rural Mallards, Anas platyrhynchos. Journal of Urban Ecology 6, 1–11. 10.1093/jue/juaa011

Kalnay, E., Cai, M., 2003. Impact of urbanization and land-use change on climate. Nature 423, 528–531. 10.1038/nature01675

Kopij, G., 2020. Distribution and numbers of waterbird species breeding in the city of Wrocław. Acta Musei Sil Sci Natur 69, 175–180. 10.2478/cszma-2020-0013

Loss, S.R., Marra, P.P., 2017. Population impacts of free-ranging domestic cats on mainland vertebrates. Front Ecol Environ 15, 502–509. 10.1002/fee.1633

Manikowska-Slepowrońska, B., Meissner, W., 2022. Factors affecting apparent survival and resighting probability of wintering mallards Anas platyrhynchos. Ornis Fenn 00–00. 10.51812/of.121697

Meissner, W., Dynowska, M., Góralska, K., Rzyska, H., 2015a. Mallards (Anas platyrhynchos) staying in urban environments have higher levels of microfungi biota diversity than do birds from non-urban areas. Fungal Ecol 17, 164–169. 10.1016/j.funeco.2015.07.004

Meissner, W., Markowska, K., 2009. Influence of low temperatures on behaviour of mallards (Anas platyrhynchos L.). Pol J Ecol 57, 799–803.

Meissner, W., Rowiński, P., Polakowski, M., Wilniewczyc, P., Marchowski, D., 2015b. Impact of temperature on the number of mallards, Anas platyrhynchos, wintering in cities. North West J Zool 11, 213–218.

Meissner, W., Witkowska, M., 2023. The effect of the temperature on local differences in the sex ratio of Mallards Anas platyrhynchos wintering in an urban habitat. Acta Oecologica 119. 10.1016/j.actao.2023.103900

Pattenden, R.K., Boag, D.A., 1989. Skewed sex ratio in a northern wintering population of mallards. Can J Zool 67, 1084–1087. 10.1139/z89-152

Pavisse, R., Vangeluwe, D., Clergeau, P., 2019. Domestic Cat Predation on Garden Birds: An Analysis from European Ringing Programmes. Ardea 107, 103–109. 10.5253/arde.v107i1.a6

Plummer, K.E., Risely, K., Toms, M.P., Siriwardena, G.M., 2019. The composition of British bird communities is associated with long-term garden bird feeding. Nat Commun 10. 10.1038/s41467-019-10111-5

Polakowski, M., Skierczyński, M., Broniszewska, M., 2010. Effect of urbanization and feeding intensity on the distribution of wintering Mallards Anas platyrhynchos in NE Poland. Ornis Svec 20, 76–80. 10.34080/os.v20.22636

R Core Team, 2021. R: A language and environment for statistical computing.

Sauter, A., Korner-Nievergelt, F., Jenni, L., 2010. Evidence of climate change effects on within-winter movements of European Mallards Anas platyrhynchos. Ibis 152, 600–609. 10.1111/j.1474-919X.2010.01028.x

Schummer, M.L., Kaminski, R.M., Raedeke, A.H., Graber, D.A., 2010. Weather-Related Indices of Autumn–Winter Dabbling Duck Abundance in Middle North America. Journal of Wildlife Management 74, 94–101. 10.2193/2008-524

Slabbekoorn, H., Ripmeester, E.A.P., 2008. Birdsong and anthropogenic noise: Implications and applications for conservation. mMol Ecol 17, 72–83. 10.1111/j.1365-294X.2007.03487.x

Swanson, D.L., Olmstead, K.L., 1999. Evidence for a proximate influence of winter temperature on metabolism in passerine birds. Physiological and Biochemical Zoology 72, 566–575. 10.1086/316696

Tryjanowski, P., Sparks, T.H., Biaduń, W., Brauze, T., Hetmański, T., Martyka, R., Skórka, P., Indykiewicz, P., Myczko, Ł., Kunysz, P., Kawa, P., Czyz, S., Czechowski, P., Polakowski, M., Zduniak, P., Jerzak, L., Janiszewski, T., Goławski, A., Dudu, L., Nowakowski, J.J., Wuczyński, A., Wysocki, D., 2015. Winter bird assemblages in rural and urban environments: A national survey. PLoS One 10, 1–25. 10.1371/journal.pone.0130299

Wang, W., Wu, T., Li, Y., Xie, S., Han, B., Zheng, H., Ouyang, Z., 2020. Urbanization impacts on natural habitat and ecosystem services in the Guangdong-Hong Kong-Macao “Megacity.” Sustainability (Switzerland) 12. 10.3390/su12166675

Wood, A.S., Scheipl, F., Wood, M.S., 2020. Package ‘gamm4.’

Zhu, J., Ding, N., Li, D., Sun, W., Xie, Y., Wang, X., 2020. Spatiotemporal analysis of the nonlinear negative relationship between urbanization and habitat quality in metropolitan areas. Sustainability (Switzerland) 12. 10.3390/su12020669

