## Supplementary Materials for "The intensity of supplementary feeding in an urban environment impacts overwintering Mallards *Anas platyrchynhos* as wintering conditions get harsher"

[4] – Regional Directorate for Environmental Protection, Chmielna 54/57, 80-748 Gdańsk, Poland

Keywords: waterbird, ecology of cities, urban ecology, duck, anthropogenic food

Table 1S. The exact date of conducting a particular counting in each of the studied winter seasons, together with the mean ambient temperature established for a given date. The temperature values below 0 °C were bolded, '- ' indicates countings omitted in the first, pilot season of the study.

| Season | | Counting occasion | | | | | | |
|---|---|---|---|---|---|---|---|---|
| | | 1st | 2nd | 3rd | 4th | 5th | 6th | 7th |
| 2017/2018 | Date | - | - | 20th Jan | 3rd Feb | 17th Feb | 4th Mar | 20th Mar |
| | Temp. | - | - | 0 °C | 3.5 °C | **-4 °C** | **- 10 °C** | **-3.5 °C** |
| 2018/2019 | Date | 22nd Dec | 5th Jan | 19th Jan | 2nd Feb | 16th Feb | 2nd Mar | 16th Mar |
| | Temp. | 0 °C | 1 °C | 1 °C | **-1 °C** | 3.5 °C | 3 °C | 5 °C |
| 2019/2020 | Date | 21st Dec | 4th Jan | 18th Jan | 31st Jan | 14th Feb | 28th Feb | 14th Mar |
| | Temp. | 5.5 °C | 2.5 °C | 4.5 °C | 3 °C | 2.5 °C | 2 °C | 4 °C |
| 2020/2021 | Date | 19th Dec | 2nd Jan | 17th Jan | 31st Jan | 13th Feb | 28th Feb | 13th Mar |
| | Temp. | 4 °C | 0 °C | **-3.5 °C** | 0 °C | 0 °C | 6 °C | 2 °C |
| 2021/2022 | Date | 18th Dec | 2nd Jan | 15th Jan | 29th Jan | 13th Feb | 26th Feb | 13th Mar |
| | Temp. | 4 °C | 1 °C | 1 °C | 2.5 °C | 2.5 °C | 3 °C | 1 °C |
